# Confidence-controlled Hebbian learning efficiently extracts category membership from stimuli encoded in view of a categorization task

**DOI:** 10.1101/2020.08.06.239533

**Authors:** Kevin Berlemont, Jean-Pierre Nadal

**Affiliations:** Laboratoire de Physique de l’Ecole Normale Supérieure, CNRS, ENS, PSL University, Sorbonne Université, Université de Paris, Paris, France; Center for Neural Science, New York University, New York, United States; Centre d’Analyse et de Mathématique Sociales, École des Hautes Études en Sciences Sociales, CNRS, Paris, France

## Abstract

In experiments on perceptual decision-making, individuals learn a categorization task through trial-and-error protocols. We explore the capacity of a decision-making attractor network to learn a categorization task through reward-based, Hebbian type, modifications of the weights incoming from the stimulus encoding layer. For the latter, we assume a standard layer of a large number of stimulus specific neurons. Within the general framework of Hebbian learning, authors have hypothesized that the learning rate is modulated by the reward at each trial. Surprisingly, we find that, when the coding layer has been optimized in view of the categorization task, such reward-modulated Hebbian learning (RMHL) fails to extract efficiently the category membership. In a previous work we showed that the attractor neural networks nonlinear dynamics accounts for behavioral confidence in sequences of decision trials. Taking advantage of these findings, we propose that learning is controlled by confidence, as computed from the neural activity of the decision-making attractor network. Here we show that this confidence-controlled, reward-based, Hebbian learning efficiently extracts categorical information from the optimized coding layer. The proposed learning rule is local, and, in contrast to RMHL, does not require to store the average rewards obtained on previous trials. In addition, we find that the confidence-controlled learning rule achieves near optimal performance.

---

As categorization occurs constantly in everyday life, the ability to learn new categories is essential to achieve accurate behaviors [Ashby and Maddox, 2005]. In laboratory experiments, associations between the continuum of sensory stimuli and the discrete categories are learned through trial-and-error protocols with simple perceptual decision tasks – see e.g. [Ghose et al., 2002, Yang and Maunsell, 2004, Freedman and Assad, 2006].

Based on experimental data in both neuroscience and experimental psychology, the modelling of category learning addresses various issues. First, it aims at elucidating the neural basis of decision making accounting for behavioral data, such as reaction times and performance, and for the observed neural activity in putative decision areas - ramping activities that can be understood as representing the accumulation of evidence [Gold and Shadlen, 2007]. From a behavioral point of view, many effects are observed during such experiments which are specific to category learning [Drugowitsch et al., 2019] - such as speed/accuracy compromise, sequential effects [Drugowitsch et al., 2019]. In the last decade, researchers have also studied the neural basis of confidence in one’s decision [Kiani and Shadlen, 2009, Jaramillo et al., 2019, Wei and Wang, 2015], and how to model the learning task within the frameworks of reinforcement learning and Hebbian plasticity [Engel et al., 2015]. In the modelling studies, authors either only consider the decision layer, with e.g. competing units associated with the different categories, or consider both the decision layer and the stimulus encoding area, with a large number of stimulus specific cells. In the present paper we are concerned with the coherent interplay between different components of perceptual decision making: stimulus encoding, neural decision making dynamics, reward-based adaptation of the coupling between the coding and decision layers. Taking advantage of experimental and theoretical results on categorical perception, here we show how and why confidence can play an important role in the course of learning.

For the modeling of behavioral performances in perceptual decision tasks, the process of accumulation of evidence is most often formalized with the Drift-to-Bound decision models (DDM) [Ratcliff, 1978, Bogacz et al., 2006, Gold and Shadlen, 2007]. The DDMs nicely account for a large variety of experimental data, are amenable to analytical analysis and fit well within a Bayesian inference framework. However, within a same sequence of trials, different conditions (such as post-error and post-correct trials [Dutilh et al., 2012]) must be fitted with different parameters values, without explicitation of the neural mechanisms controlling such changes of parameters. As a more biophysical approach, models of attractor neural decision networks [Wong and Wang, 2006] have been shown to account as well for various behavioral data such as reaction times, accuracy, as well as confidence [Wei and Wang, 2015, Berlemont et al., 2020] and sequential effects [Berlemont and Nadal, 2019], but without ad hoc changes of parameters as mentioned above in the case with DDMs. This results from intrinsic nonlinear features of the neural dynamics, which are not taken into account within the DDM framework. The cost to pay for the modeller is a higher complexity in model analysis and in the methods needed to calibrate the model against empirical data. In the following we will adopt this more biophysical modelling approach.

When taking into account the nature of the stimulus encoding, models consider a typical architecture with a coding layer of a large number of stimulus specific cells [Tajima et al., 2016, Engel et al., 2015]. The tuning curves may be chosen to uniformly cover the stimulus space, or to be adapted according to some criterion. One possibility consists in assuming that the encoding is optimized for the estimation or reconstruction of the stimulus itself [Ma et al., 2006]. However, if one considers that the optimisation is done in view of the categorization task, experimental [Goldstone, 1994, Sigala and Logothetis, 2002, Harnad, 2003, Xin et al., 2019] and theoretical analysis [Bonnasse-Gahot and Nadal, 2008, 2012] show that tuning curves of coding neurons are sharper near to than away from the category boundaries in stimulus space. These findings account for the various effects characterizing categorical perception [Harnad, 2003, Bonnasse-Gahot and Nadal, 2008].

Various experiments [Summerfield and De Lange, 2014, Meyniel and Dehaene, 2017, Meyniel, 2019] show that confidence plays a role during learning. For instance, in the context of uncertain environment, authors have shown that the sense of confidence in the stability of the environment affects the speed of learning. However, this already corresponds to a rather high level of metacognition. In order to elucidate the neural basis of confidence, researchers have focused on simplest experimental protocols, allowing to combine experimental and theoretical approaches (see e.g. [Ott et al., 2019]). Within the framework of drift-diffusion models and taking a Bayesian viewpoint, Drugowitsch et al. [2019] have shown that the optimal learning rate for categorization tasks should depend on the confidence in one’s decision, where confidence is defined as the probability of having answered correctly. Within the attractor neural network framework, confidence in one’s decision is well accounted for by the difference, at the time of decision, between the neural activities of the decision-specific neural pools [Wei and Wang, 2015, Berlemont et al., 2020]. This measure of the level of confidence is thus available at the neural, local, level, and can be used as a feedback for future decisions and for learning, in the spirit of the just mentioned statistical inference approach.

Within the framework of reinforcement learning [Sutton and Barto, 2018], different learning algorithms have been proposed to account for the learning of perceptual decision tasks. As a bio-physically plausible Hebbian learning rule [Hebb, 1949, Miller and MacKay, 1994] modulated by a reward signal [Schultz et al., 1997, Loewenstein and Seung, 2006, Gerstner et al., 2018], Reward-modulated Hebbian learning (RMHL) [Legenstein et al., 2008, 2010] has been proposed and used in various contexts. RMHL has in particular be used in models of categorical perception to adapt the weights between the encoding and the decision layers [Engel et al., 2015, Min et al., 2020]. However, such rule requires to keep track of the rewards associated to the previous trials to generate the prediction error. Moreover, the impact of the optimization of the coding layer on the efficiency of such Hebbian type learning has not been specifically addressed.

In the present work, we consider a network composed of two layers, the stimulus encoding layer and the decision layer. We study the efficiency of Hebbian learning rules acting on the weights between the two layer. We do so comparing two possible distributions of tuning curves for the coding neurons: uniformly distributed or optimized in view of the categorization task - en passant, we derive an analytical formula giving the optimal tuning curve distribution in the present context.

As learning rule, we first consider the simplest Reward modulated Hebbian rule (RMHL). We find that RMHL is not able to successfully extract categorical information from an optimized coding layer. Analyzing the failure of this rule, we propose a reward-based Hebbian learning rule controlled by confidence, the latter being extracted from the neural decision dynamics. In the proposed rule, confidence acts as a modulator of the Hebbian synaptic plasticity taking place between the coding and decision layers. The rule corresponds to a particular case of the three factor learning rule introduced by Frémaux and Gerstner [2016] as a general framework for taking into account the effect of neuromodulators. The first two factors correspond to the standard pre- and post-synaptic neural activities, and the third factor quantifies the effect of a neuromodulator. Neuromodulators may convey information about reward [Schultz, 1998], novelty [Ranganath and Rainer, 2003] uncertainty [Angela and Dayan, 2005] or attention [Angela and Dayan, 2005]. In our case, the third factor represents the effect of confidence.

When learning is controlled by confidence, we show that a network with coding neurons optimized for the categorization task leads to better performance than a network with uniform tuning curves. Moreover, we find that our confidence-controlled Hebbian learning leads to near optimal performances. Our findings show that confidence, as measured at the neural level, allows to successfully learn a categorization task with a local Hebbian rule, without keeping track of prediction errors.

## Results

### Neural circuit model and learning rules

We consider a neural circuit trained to perform a categorization task, more precisely a two-alternative-forced choice (2AFC) task. Stimuli are sampled among two categories (such as motion directions for example) and sequentially presented to the network. The categories overlap, that is the category membership of a stimulus is more or less ambiguous depending on how far it is from the class boundary in the stimulus space (*x* = 0.5). The behavioral task is to learn to distinguish the category of the stimuli through trial and error. The network is composed of two layers (Fig. 1.A), a stimulus encoding layer feeding a decision-making attractor network.

**Figure 1:**
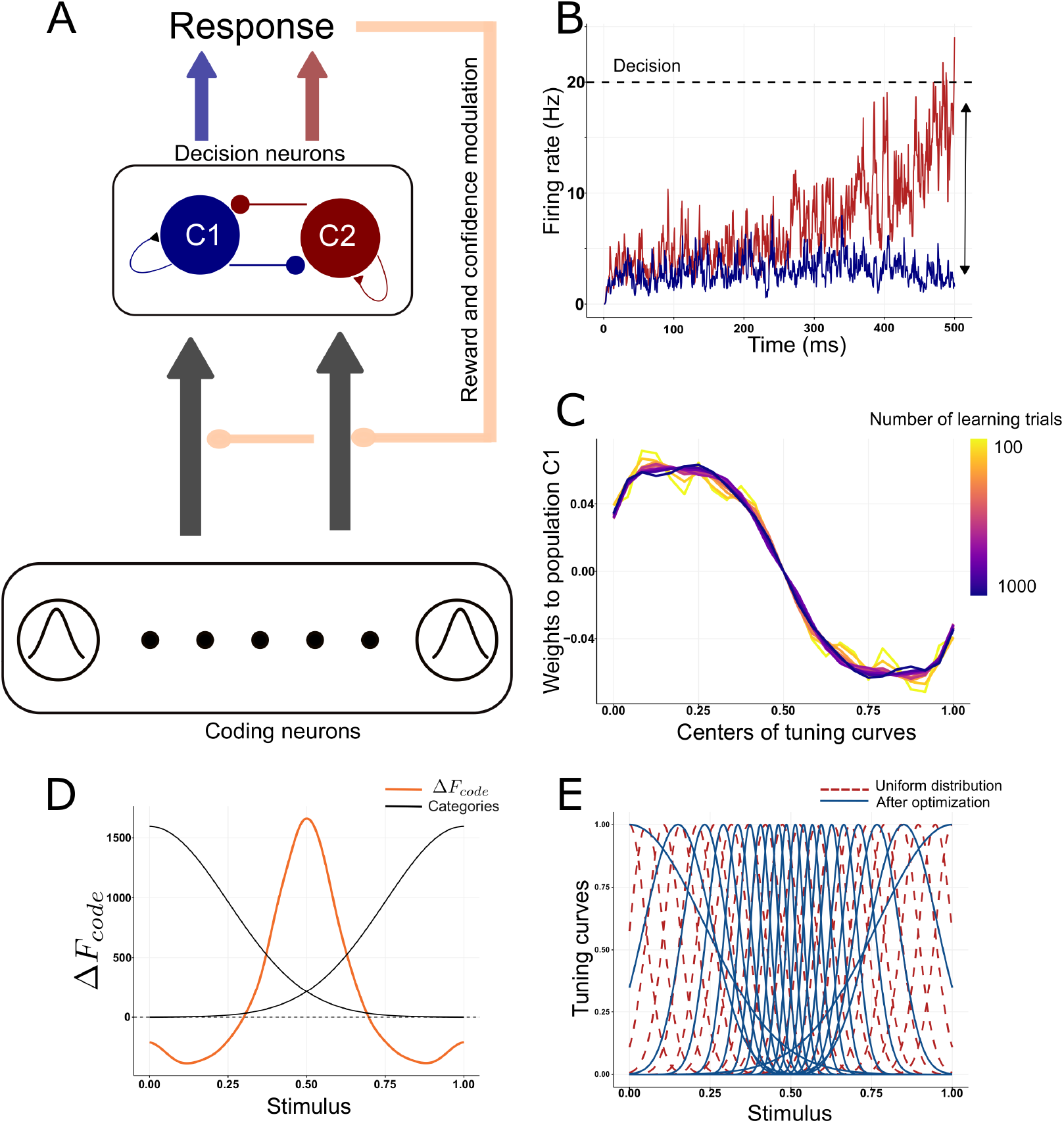
Neural circuit model. (A): Schematic of the circuit model. The network is composed of two layers. Neurons in the coding layer are tuned to specific values of the 1-d stimulus. The decision layer pools activity of the coding neurons through feedforward synapses. The category decision is obtained through competitive attractor dynamics. The synaptic connections between the two layers undergo Hebbian plasticity modulated by a reward signal. (B): Example of the dynamics of the decision layer n presentation of a stimulus. Each color represent the activity of the corresponding population. The decision is reached when the activity of one of the populations reach the decision threshold θ (dotted line). The behavioral confidence is a monotonous function of the difference Δr in firing rates between the two populations, computed at the time of decision. (C): Synaptic weights from the coding layer to one of the two neural populations, C_1_. Each color represents a different epoch (each of 100 trials) in the learning process, from 100 to 1000 trials. The coding layer is composed of 20 neurons, uniformly distributed. (D): Fisher information of the coding layer. Orange curve: difference in Fisher information between the case of an optimized layer and the one of a uniform distribution of tuning curves (see Materials and Methods). Black curves: categories used for this example. (E): Tuning curves. Red curves: tuning curves for an uniform distribution. Blue curves: tuning curves of the task-optimized coding layer. For this example, coding layer of 20 neurons. For the numerical examples (B, C, D, E), the Gaussian categories are centered at 0 and 1 respectively, and of same width α = 0.25 (black curves, panel D).

Coding neurons are modeled as Poisson neurons with stimulus-specific bell-shaped tuning curves (Fig. 1.E). We restrict to a one dimensional stimulus space, a direction that discriminates the two categories. The activity of the coding layer is pooled from the decision layer. The decision layer is composed of an attractor network of two populations (*C*_1_ and *C*_2_) that compete with each other (Fig. 1.A). A decision is reached when the activity of one population crosses a threshold *θ*. This two-units model is a mean-field approximation of a large network of spiking neurons [Wang, 2002].

In the following, we compare two different networks: one where the tuning curves in the coding layer are uniformly distributed along the stimulus axis with identical widths *w*_*i*_ = *w* (in the following referred to as the uniform network), and one where the distribution of tuning curves has been optimized with respect to the categorization task (in the following referred to as the task-optimized network. In the latter case, tuning curves are sharper near the boundary between categories [Bonnasse-Gahot and Nadal, 2008], the coding layer allocating more resources where the task is more difficult (greater ambiguity of the stimulus) - see Fig. 1.E.

For a given coding layer (optimized or not), we consider the learning occurring between the coding and decision layers through the adaptation of the synaptic connections. To analyse in detail the learning between the coding and decision layers, we do not consider the simultaneous adaptation of the tuning curves in the coding layer.

At each trial, the strength 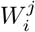 of the synaptic connection between a coding neuron *i* and the winning population (*C*_*j*_) of the trial is updated. We consider three reward-based Hebbian learning rules - which can be seen as examples of ‘NeoHebbian Three-Factor Learning Rules’ [Gerstner et al., 2018]. As reference, the first rule is a pure Hebbian reinforcement learning rule (hereafter PHL), equivalent to a Hebbian supervised learning rule:

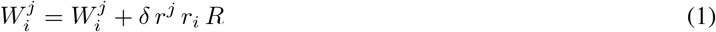

Here and in the following, *R* is the reward of the trial (1 if the decision was correct, −1 otherwise), *r*_*i*_ the firing rate of the presynaptic neuron, *r*^*j*^ the one of the post-synaptic unit, and we denote by *δ* the learning rate. The second rule is the Reward Modulated Hebbian Learning rule (RMHL),

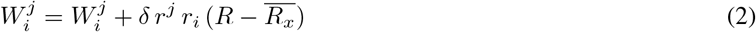

where 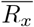 is the moving average of the reward obtained on a short recent time window for the particular stimulus *x* on that trial. Note that 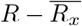, if non-zero, has the sign of *R*. Finally, the confidence-controlled reward-based Hebbian learning rule (CCHL) that we introduce in the present paper is defined by:

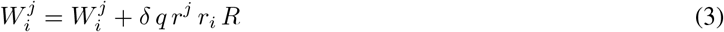

where *q* is the confidence control parameter defined by:

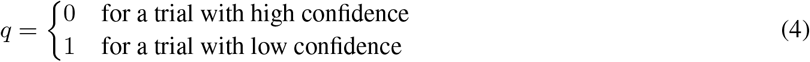

That is, there is no updating of the weights for high confidence trials, and the learning rule is the pure reward-based Hebbian one in case of low confidence.

Following [Wei and Wang, 2015, Berlemont et al., 2019], confidence is here modelled as a function of the difference Δ*r* between the neural activities of the two populations of the decision layer at the time of decision. Jaramillo et al. [2019] have proposed a pulvinar-cortical circuit model that can extract this quantity. In a previous work, we have shown that this difference in neural activity provides a neural basis for behavioral confidence in human [Berlemont et al., 2019]. We consider that a trial corresponds to a high (resp. low) confidence trial when the difference in activity Δ*r* at the time of the decision is greater (resp. lower) than a threshold level *z*.

More details about the model, and a discussion about the link between CCHL algorithm and machine learning algorithms, can be found in the STAR methods section.

### Learning: Reward-Modulated Hebbian learning

First, we study the simplest case of pure reward-based Hebbian learning. Before learning, the synaptic connections are initialized with random values leading to performance at chance level. Fig. 1.C represents the weights of the coding neurons towards the neural population *C*_1_ during learning.

In Fig. 2.A we compare the performances achieved by the network for different sizes, for the two types of coding layers (uniform and task-optimized tuning curves) after 2000 trials. The large number of trials is chosen (here and for most results shown in this paper) so that the weights stabilize, allowing to obtain well-defined average performances for all values of parameters. However, it is important to note that the network does learn to perform the behavioral task with a rather good success rate, and this after only a small number of trials as can be seen in Fig. 2.C and D.

**Figure 2:**
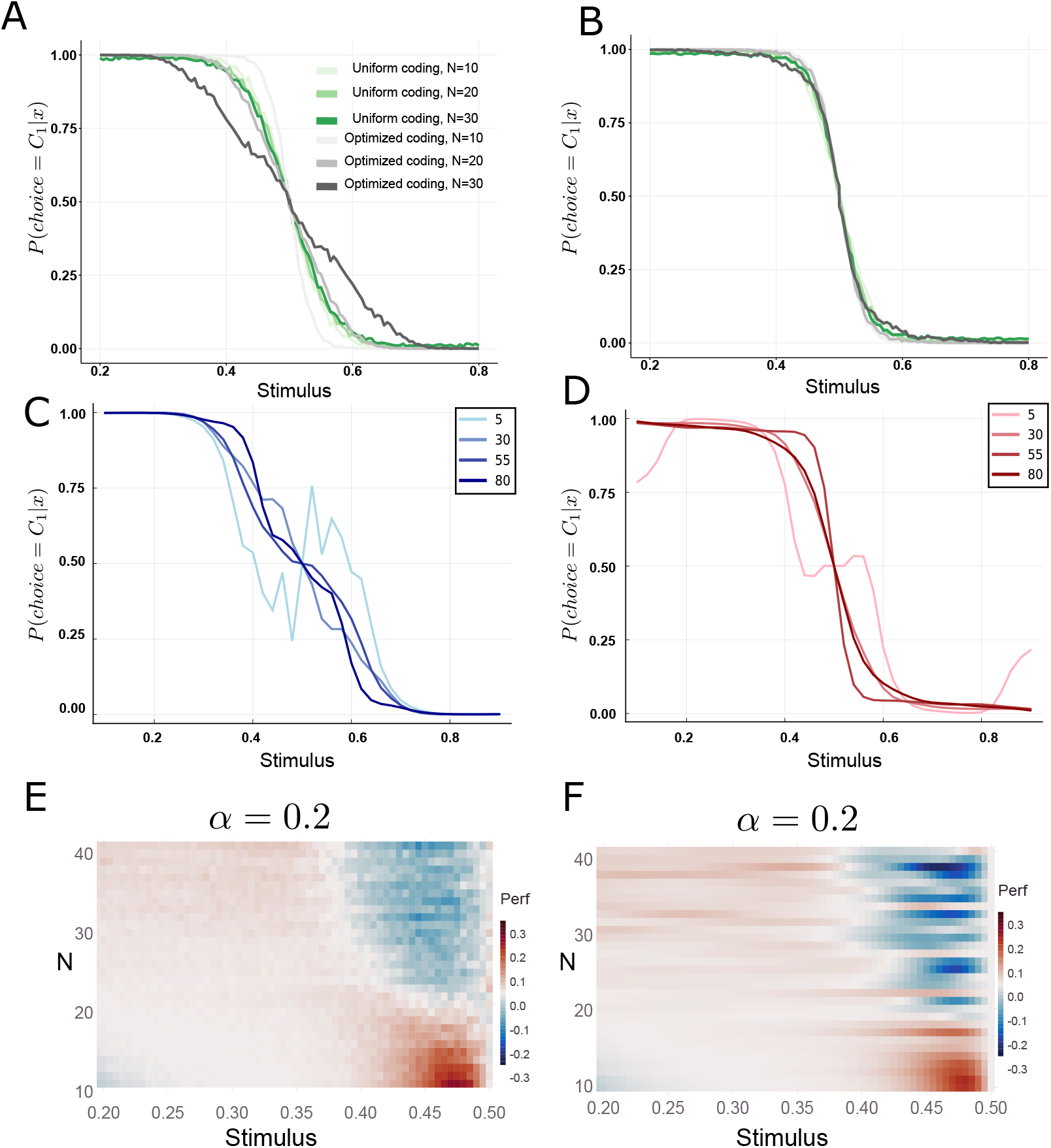
Performance of the network after learning: Probability of choosing category C_1_ with respect to the presented stimulus x after 2000 trials. (A) and (B): Effect of network size. (A): Case of the PHL rule. (B) case of the CCHL rule. For both panels, Green curves: uniform network; Grey curves: task-optimized network. The number of neurons in the coding layer is indicated by going from light to dark colors. (C) and (D): Early stages of learning illustrated on the case of PHL. The gradient of color represents different snapshots of the network state after n = 5, 30, 55, 80 learning trials. (C, blue curves): task-optimized coding layer; (D, red curves): uniform coding layer. The behavioral task is learned after 30 trials. All panels: The width of the Gaussian categories is α = 0.25, the number of coding cells is N = 20; the threshold for confidence control is z = 15 Hz. (E) and (F): Difference in performances after 2000 learning trials between networks with uniform and task-optimized coding layer, for the pure Hebbian learning rule (PHL, panel E) and the Reward-modulated rule (RMHL, panel F). Red (resp. Blue) corresponds to a positive (resp. negative) difference, meaning that the network with an optimized coding layer has better (resp. worse) accuracy that the one with a uniform coding layer. The x-axis gives the stimulus x (category boundary at x = 0.5), and the y-axis to the number of neurons N. The width of the categories (centered at 0 and 1) is α = 0.2.

In panels 2.E and 2.F, we show the difference in accuracy between a task-optimized network and an uniform one in the plane stimulus ambiguity/network size: Red (resp. Blue) regions correspond to higher (resp. lower) performances for the task-optimized coding layer with respect to the uniform one. We show the results for the PHL (Fig. 2.E) and the RMHL (Fig. 2.F), at medium overlap of the categories.

Surprisingly, the performance appears to be generally better for the non-optimized network. First, when the distribution of the tuning curves is uniform, the performances do not vary significantly with *N*, whereas, for a task-optimized network, the accuracy *decreases* with the number of coding neurons. Secondly, when the number of neurons increases, a network with a non-optimized coding layer performs better that a task-optimized network. It is only for small values of *N* near the category boundary that the performances are better with an optimized coding layer (2.E and 2.F). The region where uniform coding performs better increases with *α*. However, the difference in accuracy between the two networks decreases if one keeps increasing *α* (see Supplementary Fig. 2 for an example). This is explained by the fact that, when the categories widths are wider, the overlap between the tuning curves is larger, and the optimal code becomes more similar to an uniform one.

These apparent paradoxical behaviors all result from the same reason. In the case of the task-optimized coding layer, the tuning curves are sharp at the boundary between categories in order to maximize the Fisher information. This means that a neuron close to the boundary will only emit spikes for stimuli close to the center of its tuning curve. Thus, the strength of the associated synaptic connections will less often be updated than for a neuron far away from the boundary, whose firing rate is higher in average. Finally, the weights tend to decrease to zero close to the category boundary, in effect ‘fighting’ against the optimal tuning of coding cells (Fisher information should be large at such location). Hence, during learning the network progressively loses information as finely tuned coding neurons tend to be associated with a synaptic connection of decreasing strength.

### Learning: Control by confidence

In order to counterbalance the loss of information near the category boundary, the idea would be to only update the synaptic connections when it is useful to do so for the network, that is when it would gain information in doing so. In other words, once the behavioral task begins to be acquired, one would like to finely tune the learning by only considering the hard trials, hence the ones near the categories boundary [Krauth and Mézard, 1987, Guyon et al., 1996, Alemi et al., 2015].

This can be achieved through the control of learning by the confidence. Let us first see how confidence, as measured by the difference Δ*r* between the activities of the decision units at the time of decision, is being built during the learning process. Fig. 3.F shows the mean level of confidence of the network, defined as the mean value of Δ*r*, with respect to the stimulus *x* that is presented. One can see that confidence builds up at the early stage of the learning process. One can state that the notion of confidence is present in the network from the very beginning of the learning process. As expected, the confidence level decreases as the stimulus is chosen closer to the categories boundary. As the number of learning steps increases, convergence in confidence level is fast far from the boundary, and slow near the boundary. As the task is learned more accurately, the representation of confidence within the attractor network becomes sharper at the boundary.

**Figure 3:**
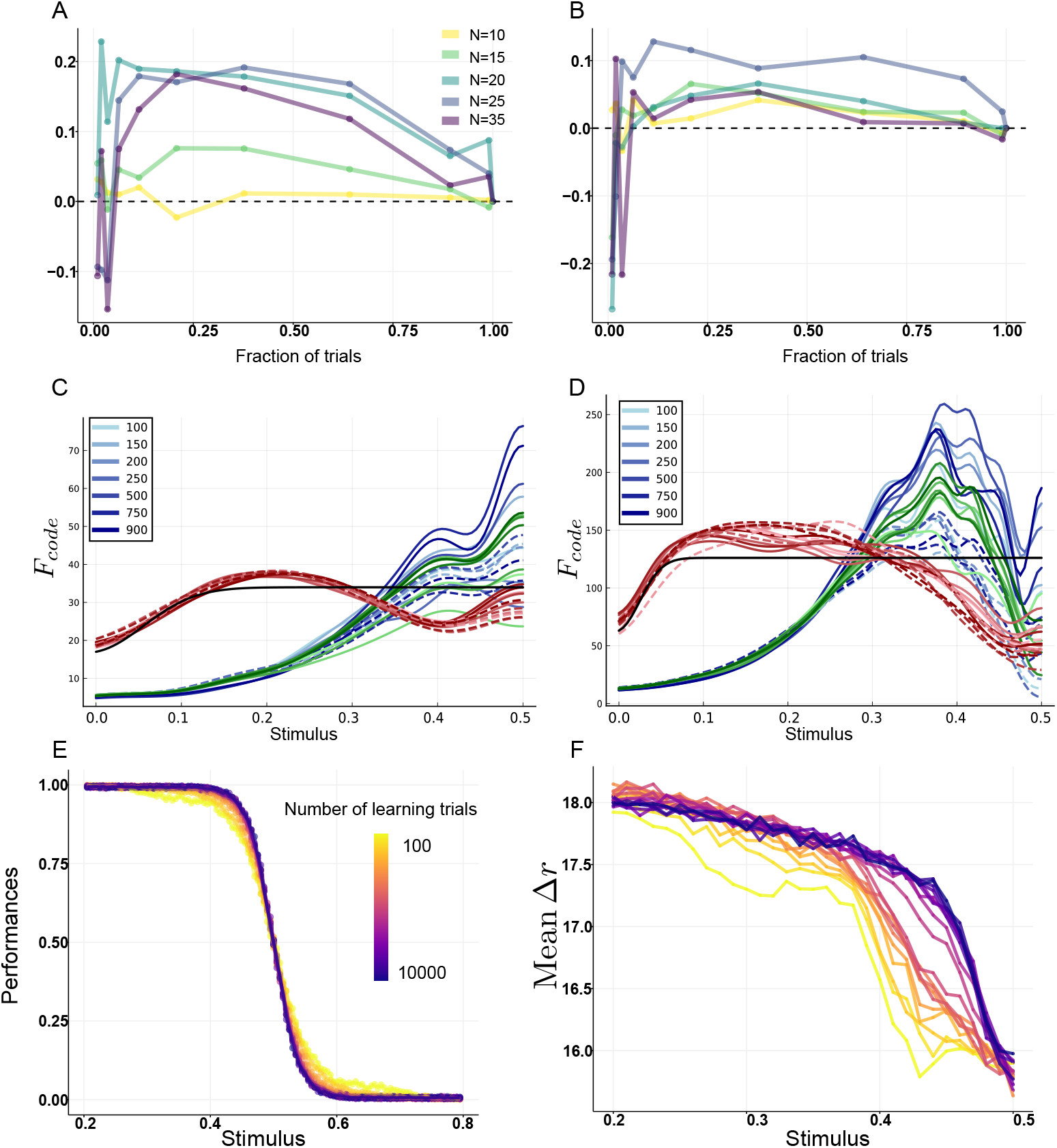
Learning with the confidence-controlled Hebbian learning rule (CCHL). (A): Effect of confidence control on the performances for a network with an optimized coding layer after 2000 trials. The y-axis represents the difference in performances, at an ambiguity x = 0.45, for the learning rule controlled by confidence and for the learning without confidence-control. The x-axis represents the percentage of trials for which Δr was lower than the threshold z. (B): Same as (A) but for a neural circuit model with an uniformly distributed coding neurons. (C) and (D): Fisher information of the coding layer taking into account the gain modulation due to the synaptic weights (see Material and Methods), for N = 15, panel (C), and N = 35, panel (D). The gradient of color represents different snapshots of the network during learning (after an increasing number of trials, from 100 to 900). The network with an uniform coding layer is represented in red and the one with an optimized coding layer in blue. The dashed lines stands for the pure Hebbian learning rule; and the plain lines for the confidence-controlled Hebbian rule. The green color corresponds to the optimized network with the RMHL rule. (E): Performance for the task-optimized network for different numbers of learning trials (logarithmic scale). (F): Δr averaged over 1000 trials with respect to the stimulus ambiguity. The different curves correspond to different numbers of learning steps. The number of coding cells is N = 20 for panels E and F. All panels: The width of the Gaussian categories is α = 0.25. Panels (C) to (F), the threshold for confidence control is z = 15 Hz.

We thus expect the CCHL to make use of this sharpening representation of confidence during learning to improve the performances of the network near the categories boundary.

In Fig. 3.A and B we consider the accuracy near the boundary (stimulus *x* = 0.45) for the confidence-controlled learning rule as compared to the pure Hebbian one. The important parameter in the CCHL scheme is the confidence threshold *z*. This parameter determines the fraction of trials that will be considered as *low* or *high* confidence trials. The *x*-axis in Fig. 3.A and B represents the fraction of low confidence trials as one varies the threshold *z*. We observe that, for low values of *N*, the confidence control does not have any impact on the performances. This is easily understood: due to the low amount of resources, the network can not have really sharp tuning curves at the boundary and is less affected by the decay of weights near the boundary. However, as soon as one increases the number of neurons *N*, confidence control has an impact on the performances. The tendency of all the curves is the same: an increase of performance when the confidence threshold decreases, until it reaches a maximum at a confidence threshold corresponding to *∼* 25% of low-confidence trials. Finally, as one keeps decreasing the threshold *z*, the performances decrease: learning becomes inefficient as there are too few stimuli leading to a change in weight. Finally, we find that the value of confidence threshold *z* leading to the best performances is stable with respect to the number of trials (Supplementary Fig. 1), with a plateau around *∼*25% of low-confidence trials. These results imply in particular that the control by confidence can be kept constant during learning and does not need to adapt to the current state of learning.

In Fig. 3.C and D we represent the Fisher information (*F*_*code*_) of the coding layer combined with the synaptic weights (see Material and Methods), through the learning process. We note that the impact of confidence depends strongly on the coding layer. For an uniform coding, the Fisher information is almost unchanged after learning, at low or high *N*. For an optimized coding layer one can observe a strong modification of the Fisher information during learning. The confidence control leads to an increase of *F*_*code*_ near the boundary by giving more weights to the neurons closer to the categories boundary. CCHL thus efficiently makes use of the optimized coding layer to increase the performances where the task is more difficult.

In Fig. 3.E-F, we represent the performances and the mean confidence with respect to the presented stimulus through learning (category boundary at *x* = 0.5). We show the results starting at 100 learning trials in order for the network to have already learned how to perform the task. At 100 trials near the boundary (*x ∼* 0.4−0.5), confidence is relatively flat and start to increase significantly only for *x >* 0.4. Then confidence sharpens during learning. Thus, the learning process occurs with a bootstrap effect. In a first stage, the network begins to learn the decision task. During this process, confidence starts to build up. Then the network can efficiently make use of confidence to improve its performances. This is to relate to the study of Guggenmos et al. [2016] where the authors showed that confidence can be used to improve perceptual learning in the absence of external feedback when the decision task was already learned.

### Confidence-Controlled learning efficiently extracts information from the task-Optimized coding layer

We now compare the performances for the optimized and the uniform coding layer for the confidence-controlled scheme. We present the results in Fig. 4, the analogous of Fig. 2, panels E and F, showing the differences in accuracy between a network with an optimized coding layer, and one with an uniform coding layer, in the plane stimulus ambiguity/network size, and for two values of the categories width, corresponding to medium and large overlaps of the categories (see Supplementary Fig. 3 for the late stage learning).

**Figure 4:**
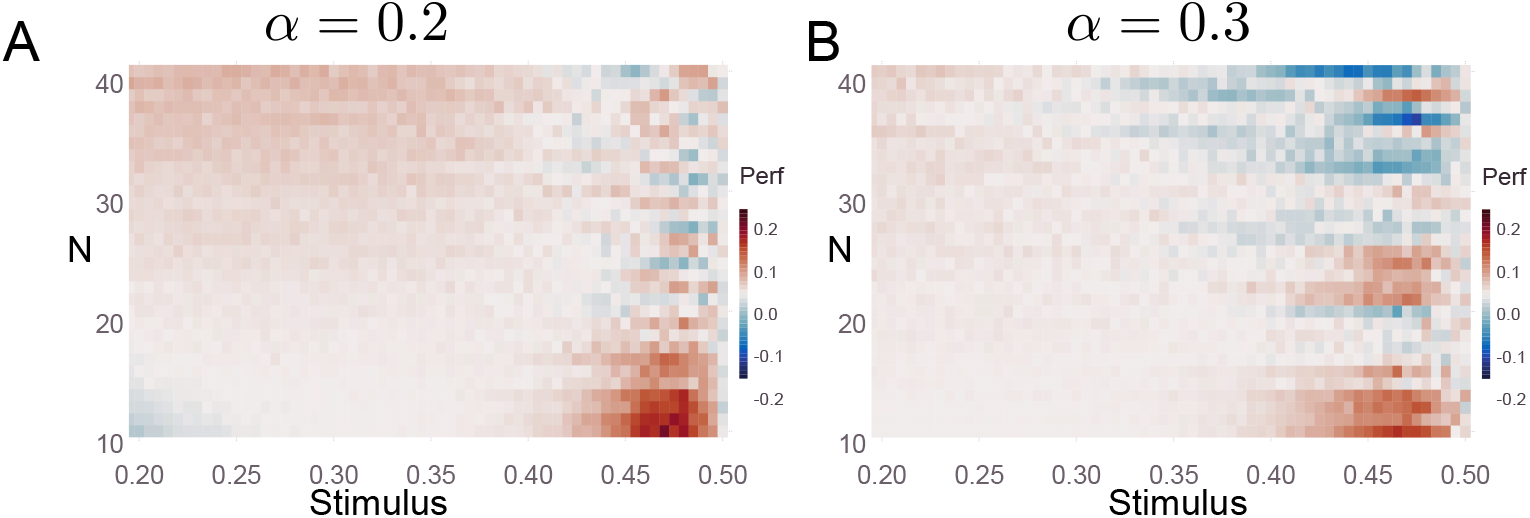
Difference in performances after learning (2000 trials, with control by confidence, threshold z = 15 Hz) between networks with task-optimized and uniform coding layer. Red (resp. Blue) corresponds to a positive (resp. negative) difference, meaning that the network with an optimized coding layer has better (resp. worse) accuracy that the one with a uniform coding layer. The x-axis gives the stimulus x (category boundary at x = 0.5), and the y-axis to the number of neurons N. The width of the categories (centered at 0 and 1) is α = 0.2 for (A) and α = 0.3 for (B).

We note that with the optimized coding layer the performances are better, independently of the number of neurons *N*. Indeed, by focusing on trials with low confidence, the network tends to decrease the synaptic weights far from the boundary and increase the ones close to the boundary (Supplementary Fig. 4) With higher weights close to the category boundary, the decision network takes advantage of the sharp tuning curves in the optimized coding layer to obtain a better accuracy.

In Supplementary Fig. 5, we compare the performances of the confidence-controlled learning rule with the ones of the RMHL. In agreement with the results on the Fisher information, we find that confidence-controlled Hebbian learning outperforms this learning scheme which requires to store the mean reward obtained so far for this stimulus.

### Confidence-controlled Hebbian rule almost achieve optimal performances

We ask now how the performances obtained with the confidence-controlled rule compare to the optimal ones that such network could achieve on average over the stimulus space, regardless of the learning rule. The way we define and compute the optimal performances is given in the Material and Methods section. Fig. 5 presents the difference between the optimal performances and the ones from the confidence-controlled Hebbian learning rule, at different stages of the learning process for a category width *α* = 0.2, and for both the non-optimized and the optimized coding layer. We first note that near the boundary, the network with optimized coding layer has better performances than the optimal model. However, the global performances are lower because it performs worse when the stimuli are a bit further from the boundary. Surprisingly, if we increase the number of trials (Fig. 5.C and D), the performances of the neural circuit model with a uniform coding layer do not increase. For the optimized coding layer, the accuracy increases and the network performances becomes very close to the optimal ones, especially at small network sizes. In Supplementary Fig. 6 we show that these qualitative results are observed at different values of the categories width.

**Figure 5:**
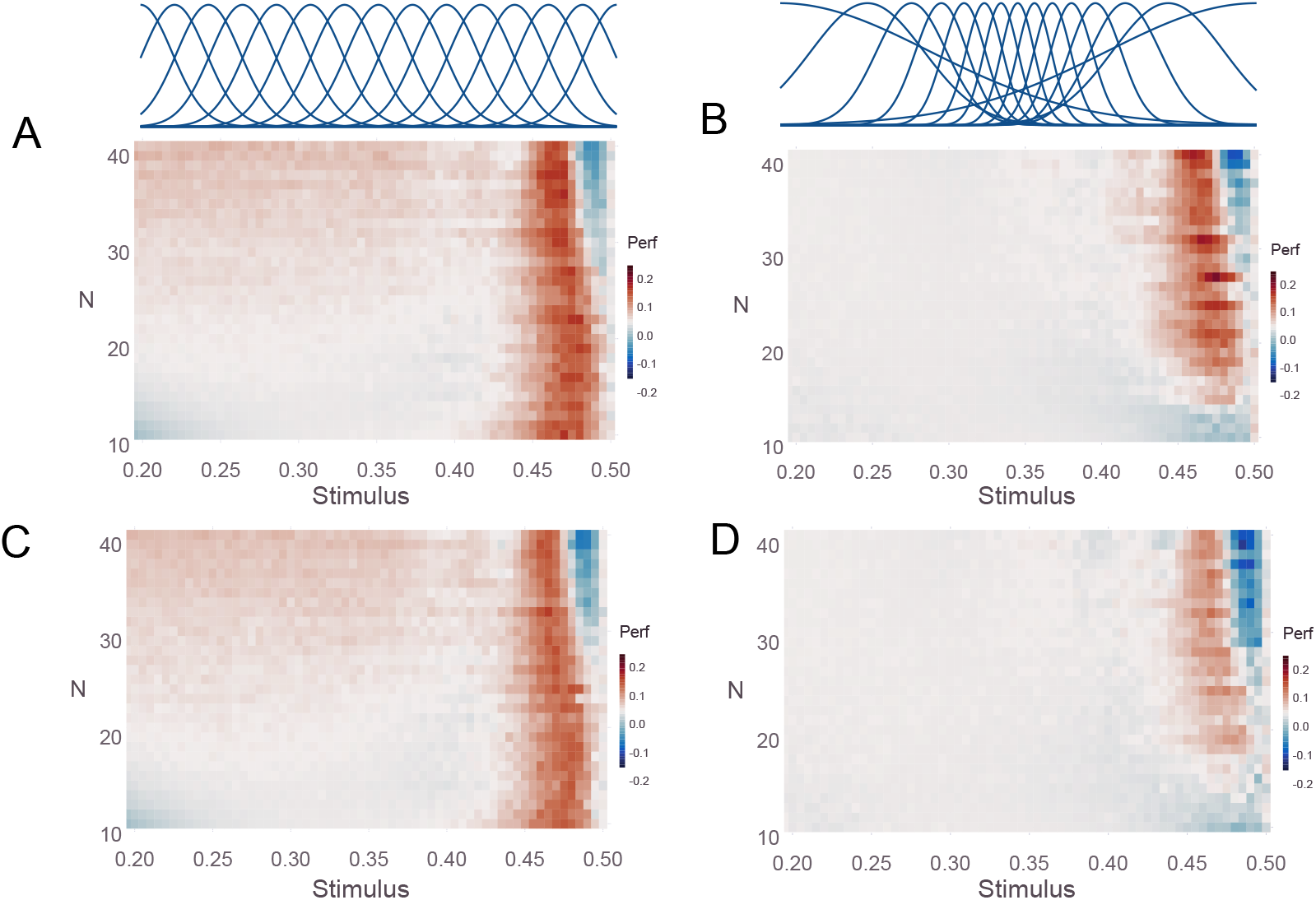
Difference between confidence-controlled Hebbian learning and optimal performances. (A) and (B) results for 2000 learning trials. Red (resp. Blue) corresponds to a positive (resp. negative) difference, meaning that the performances for the neural network with confidence-controlled learning are less (resp. higher) than the optimum performances for this neural architecture. (A) corresponds to the case where the coding layer is uniform, and (B) when it is optimized. (C) and (D): same as (A) and (B) respectively, but for 10000 learning trials. All panels: categories of width α = 0.2.

## Discussion

In this work, we proposed a learning mechanism for categorization tasks that do not rely on a prediction error signal. Comparing with both a standard reward-based Hebbian rule and the reward-modulated Hebbian rule, we showed that a scheme controlled by confidence increases significantly the performance of the neural circuit model for the categorization task, and actually achieves almost optimal performances.

The learning considered here is the one of the weights between a stimulus encoding layer and an attractor decision network. We investigated the impact on learning of having a task-specific encoding as compared to the case of an uniform coding of the stimulus. Our work demonstrates that reward-modulated Hebbian learning fails to extract the relevant information from an optimized coding layer, whereas the confidence-controlled scheme is on the contrary, particularly efficient in such cases. In addition, in contrast to RMHL, the proposed confidence-controlled rule does not need to store past average rewards.

The adaptation of the tuning curves within coding layers has been studied experimentally [Fritz et al., 2010, Xin et al., 2019]. Authors have shown that task-specific neural representations develop across different cortical areas [Cromer et al., 2010, Fitzgerald et al., 2011] during the learning of a perceptual task. These representations are accompanied by a modification in the neurons tuning properties. In monkeys trained to classify directions of random dot motion into two arbitrary categories, tuning changes are observed for neurons in the lateral intraparietal (LIP) area. In trained monkeys, individual neurons tend to show smaller differences in firing rate within categories, and larger differences between categories [Freedman and Assad, 2006], whereas in naive animals LIP neurons represent directions uniformly with bell-shaped tuning functions [Fanini and Assad, 2009]. Similar reorganization has also been observed in mice for an auditory categorization task [Xin et al., 2019]. Similar adaptation may also occur in an unsupervised way in the absence of a categorization task [Köver et al., 2013].

The adaptation of the coding layer has also been studied computationally [Bonnasse-Gahot and Nadal, 2008, Engel et al., 2015, Tajima et al., 2016, 2017, Min et al., 2020]. Whereas most models of decision-making consider an uniform coding of the stimulus before the decision part [Beck et al., 2008, Drugowitsch et al., 2019], few models analyse the nature of a stimulus coding layer optimized in view of a categorization task [Bonnasse-Gahot and Nadal, 2008]. In agreement with the experimental findings, theoretical predictions with a two layers feedforward architecture give that tuning curves in the coding layer are sharper at the vicinity of a decision boundary. In addition, within a purely statistical framework, a gradient-based supervised learning rule allows to learn the tuning curves parameters [Bonnasse-Gahot and Nadal, 2008]. Authors have proposed an alternative type of model, assuming a top-down modulation of the decision layer onto the coding layer [Tajima et al., 2016, 2017]. To obtain the adaptation of the tuning curves, authors use a top-down modulation under the form of a reward-modulated Hebbian learning [Engel et al., 2015, Min et al., 2020]. However, the efficiency of the resulting neural coding has not been discussed.

It will be interesting to see how, for the neural network architecture studied in the present work, both the tuning curves parameters and the weights from the coding layer to the decision attractor network can be efficiently learned altogether with confidence-controlled Hebbian learning rules.

Learning is accompanied by a sense of confidence about the different predictions [Nassar et al., 2010]. This sense of confidence plays a functional role in learning [Nassar et al., 2010, Meyniel and Dehaene, 2017] as it sets the balance between predictions and new information. Many studies report the existence of surprise signals in the brain, i.e a strong signal in presence of unexpected stimulus [Hillyard et al., 1971, Summerfield and De Lange, 2014]. Theoreticians have studied different models of surprise-based learning, in the absence of reward (see [Gerstner et al., 2018] for a review). More recently, making use of fMRI [Meyniel and Dehaene, 2017], EEG [Jepma et al., 2016, Nassar et al., 2019] or MEG [Meyniel, 2019] techniques, it has been shown that this surprise signal is controlled by confidence. Confidence has been shown to grade the reward signal and impact the subsequent learning in a categorization task with mice [Lak et al., 2020]. This control of the reward signal by confidence could be crucial to implement adjustable learning rates in the brain [Behrens et al., 2007, Meyniel and Dehaene, 2017]. We recall that, within the framework of attractor neural networks, confidence is given by a local signal intrinsic to the nonlinear neural dynamics.

From a theoretical point of view, the roles of surprise and novelty in the learning process have been studied [Faraji et al., 2018, Xu et al., 2020]. Xu et al. [2020] found that novelty and surprise play different roles in the learning process: novelty drives exploration while surprise modulates the learning rate. To investigate the impact of surprise, authors have proposed behavioral experiments where the feedback is corrupted, and thus unreliable [Varrier et al., 2019]. Such unreliable feedback leads to participants relying more strongly on prior beliefs to perform the task. In our framework, there are certain sets of parameters for which the impact of confidence is different depending on the unreliable feedback. For instance, unreliable feedback can lead to an increase in the utility of using confidence-controlled learning instead of reward-modulated learning. However, doing a more detailed analysis goes behind the scope of this paper as the optimal distribution of tuning curves strongly depends on the unreliable feedback. Still, our analysis suggests that it would be interesting to make experiments in which the level of unreliable feed-back is not uniform in stimulus space, but depends on the distance to the category boundary.

We note that our proposed learning scheme is reminiscent of machine learning heuristics selecting a subset of the available learning examples to efficiently learn a task. For instance, the informative vector machine algorithm [Herbrich et al., 2003, Lawrence and Platt, 2004] tends to choose points that are maximally informative. As a variant of the Perceptron algorithm, the minimum-overlap learning rule [Krauth and Mézard, 1987] leads to a learning with optimal margin by making use of a similar mechanism. In our framework, one can approximate the network behavior replacing the decision layer by a softmax unit (in the spirit of standard machine learning networks). Doing so, one obtains that the optimization by gradient descent leads to a learning rule with a term of the form of the confidence-controlled Hebbian learning (see STAR Methods). This means that the CCHL algorithm approximates a gradient descent algorithm making use of internal variables of the network. Hence we have shown that a biophysical learning rule can implement an efficient machine learning algorithm, making thus a link with a general issue at the interface between neuroscience and artificial intelligence [Whittington and Bogacz, 2019].

Finally, our confidence-controlled learning scheme is in the spirit of the Three-Threshold Learning Rule (3TLR) [Alemi et al., 2015] proposed for the encoding and retrieval of memories in recurrent neural networks. There, it is the underlying local field which provides to a given neuron the analogous of a confidence signal: Hebbian potentiation/depression occurs only if the local field is above/below some threshold (smaller than the one for the emission of a spike). The authors show that this 3TLR leads to a storage capacity close to the maximum theoretical capacity. Similarly, we have shown that the confidence-controlled rule achieves near-optimal decision performances.

## Acknowledgments

We are grateful to Laurent Bonnasse-Gahot for useful discussions and suggestions. KB acknowledges a fellowship from the ENS Paris-Saclay. KB acknowledges the support from the Office of Naval Research N00014-17-1-2041 (to Xiao-Jing Wang).

## STAR Methods

### Contact for reagent and resource sharing

Further requests for resources should be directed and will be fulfilled by the Lead Contact, Kevin Berlemont.

### Methods details

#### Neural circuit model and numerical protocol

The neural model is composed of two layers: the coding layer and the decision layer.

##### Categories and stimuli

We consider a one-dimensional stimulus, *x ∈* [0, 1]. Each stimulus belongs to one of two categories, *C*_1_ and *C*_2_. The categories are characterized by the probabilities *P* (*x*|*µ*), *µ* = *C*_1_, *C*_2_ taken as Gaussian in all simulations, with centers at 0 and 1, and width (std) *α*. The category boundary is thus at *x* = 0.5. The input *x* thus also measures the signal ambiguity.

##### Coding layer

The coding layer consists in *N* independent Poisson neurons whose firing rates are described by their bell-shaped tuning curves [Tolhurst et al., 1983]. For a given input *x*, the tuning curve of the coding cell *i* = 1,…, *N*, is

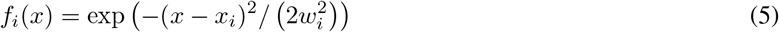

where *x*_*i*_ and *w*_*i*_ are, respectively, the center (preferred stimulus) and the width of the tuning curve.

##### Decision layer

The decision network consists in two populations *C*_1_ and *C*_2_ representing the categorical choice [Wang, 2002]. This circuit pool activity from the coding layer and produce a noisy winner-take-all dynamics. This behavior is obtained through global inhibition and recurrent excitation [Wong and Wang, 2006].

##### Sequences of trials

Each simulation consists in a sequence of trials. Each trial consists in the presentation of a randomly chosen stimulus *x* followed by a 1 s intertrial interval. During the presentation of the stimulus, the neurons in the coding layer fire a Poisson train with rate given by Eq. [5]. The stimulus presentation lasts until one of the population of the decision circuit reaches a threshold *θ* of 20 Hz. The choice made by the network on this trial corresponds to the category associated to the winning population. Once a decision has been made, there is no more input from the coding neurons to the decision neurons, while the decision neurons receive an inhibitory current (corollary discharge) [Engel et al., 2015, Berlemont et al., 2019, Berlemont and Nadal, 2019]. This brings the activity back towards the resting state, allowing the network to engage into the next trial.

##### Plasticity

The synapses connecting the coding layer to the decision layer are plastic [Schultz et al., 1997, Loewenstein and Seung, 2006]. At the end of each trial, the weights 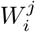 are updated according to the considered learning rule, Eq. [1], [2] or [3]. The reward is equal to *R* = 1 on correct trials and *R* = −1 on incorrect trials. The learning rate is *δ* = 0.0025. For each rule, we also added a synaptic normalization mechanism after the update of the weights (Oja’s rule), 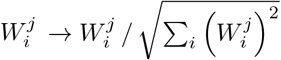. This norm-preservation mechanism prevents the divergence of the learning algorithm in the cases PHL and CCHL. It is not necessary for the RMHL but it is made here to make proper comparisons with the two other rules.

For simplicity we imposed at all times the symmetry 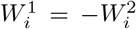 for every *i*. This is just a way to to speed up the simulations, but is in noway necessary. Note that actually the updating rule eventually leads to symmetric weights even when starting with non symmetric weights. In practice, we only apply the weight modification to the winning population, then we symetrize and normalize the weights. Finally, since at the time of decision the firing rate of the wining population is equal to the decision threshold *θ* (with small fluctuations), for each learning rule, we replace *r*^*j*^ by *θ*, here 20Hz.

Before each numerical simulation, the synapses are initialized from an uniform distribution between [−1, 1]. Unless otherwise specified, the performance of the network for a specific input is computed as the average over 2000 trials.

#### Neural circuit model - Decision layer

Here we give details on the dynamical equations of the decision layer. We model the decision layer by an attractor neural network, considering the mean-field approximation reducing the network to two effective population units Abbott and Chance [2005], Wong and Wang [2006]. Each excitatory neural population is described by a single variable *s* representing the fraction of activated N-methyl-D-aspartate receptor conductance, governed by:

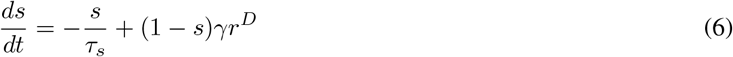

with *γ* = 0.641 and *τ*_*s*_ = 100 ms. The firing rate *r*^*D*^ of a decision unit is given by:

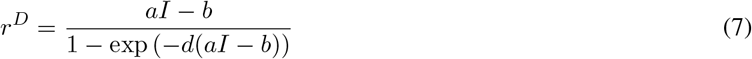

with *I* the corresponding total synaptic current, *a* = 270Hz/nA, *b* = 108Hz and *d* = 0.154s. The synaptic currents input to populations *C*_1_ and *C*_2_ are, respectively:

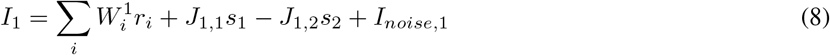

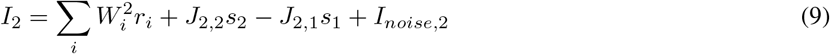

with *J*_*j,j*′_ the synaptic couplings between decision units (*J*_1,1_ = *J*_2,2_ = 0.2609 nA and *J*_1,2_ = *J*_2,1_ = 0.0497 nA). The input received by the decision unit *j* from neuron *i* in the coding layer is 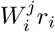, with 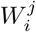 the synaptic coupling and *r*_*i*_ the spike train of the Poisson neuron *i* (resulting from the presentation of a stimulus), convoluted with a 100 ms interval. Finally, the noise terms in the synaptic currents correspond to Ornstein-Uhlenbeck processes:

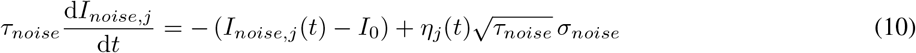

with *τ*_*noise*_ = 2 ms a synaptic time constant filtering the white-noise, and *I*_0_ = 0.3255 nA.

When the firing rate of one of the decision units reaches a decision threshold *θ*, the decision is made, and the stimulus is removed. In the simulations, we take *θ* = 20 Hz.

During a sequence of trials, after a decision has been made, in the absence of stimulus, we add a relaxation dynamics taking into account a non specific inhibitory input to each decision unit, the corollary discharge Engel et al. [2015], Berlemont et al. [2019], Berlemont and Nadal [2019]. The current of the corollary discharge is of the form:

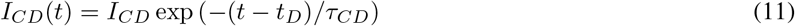

with *t*_*D*_ the time of the decision, *I*_*CD*_ = −0.05 nA and *τ*_*CD*_ = 200 ms. Under such inhibitory current, the network relaxes toward the neutral resting state, until a new stimulus is presented.

#### Optimization of the coding layer

Most works on optimal perceptual coding consider an encoding layer optimized for the estimation or reconstruction of the stimulus itself. In contrast, here we consider a neural encoding optimized in view of the categorization task. Within a Bayesian/information theoretic approach, the neural code should maximize the mutual information between the category membership and the neural activity in the coding layer [Bonnasse-Gahot and Nadal, 2008].

In the limit of a large number of coding cells, this mutual information reads

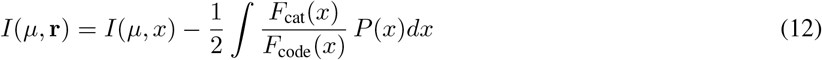

with *µ* the category membership, *x* the stimulus and **r** the neural response of the coding layer. *I*(*µ, x*) is the mutual information between category membership and stimulus, a constant characterizing the signal information content (how much the stimulus carries information about the category). *F*_code_ is the (standard) Fisher information of the neural code, measuring the sensitivity of the neural activity with respect to small changes in the stimulus value. *F*_cat_ is the categorical Fisher information quantity, a property of the signal: it measures how a small change in stimulus value affects the category likelihood. More details can be found in [Bonnasse-Gahot and Nadal, 2008, 2012].

To derive the optimal distribution of tuning curves for the coding layer, we adapt to our framework a method recently introduced for the optimal coding of a stimulus [Ganguli and Simoncelli, 2014]. Assuming that the set of tuning curves fully tiles the input space with some local density *d*(*x*), one can replace the Fisher information *F*_*code*_(*x*) by *d*(*x*)^2^. We then minimize the integral in the Eq. [12] above under the constraint of a fixed number of neurons. The loss function is then the following:

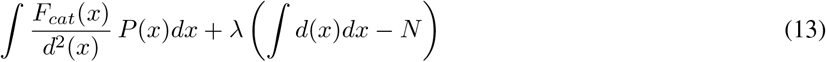

with *λ* a Lagrange parameter. Taking the derivative with respect to *d*(*x*), we obtain the optimal local density:

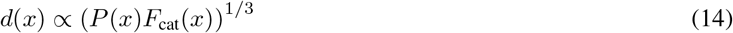

The widths of the tuning curves are given by 1*/d*(*x*). For the numerical simulations with a given number *N* of coding cells, we discretize the function 1*/d*(*x*) to get the *N* preferred stimuli.

Since the categorical Fisher information is larger where the stimulus is more ambiguous, one recovers from Eq. [14] the fact that tuning curves are sharper near the category boundaries [Bonnasse-Gahot and Nadal, 2008].

#### Fisher information quantities

We recall here the definitions of the Fisher information quantities mentioned in this paper:

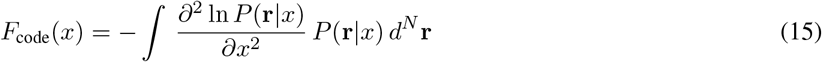

and

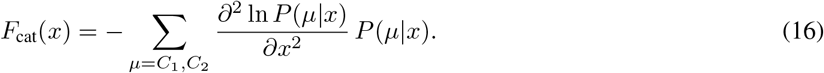

For Gaussian categories, *F*_cat_(*x*) is easily computed.

Given that the input to the decoding unit *C*_1_ from the coding cell *i* is 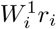, one may consider that the effective output of the coding cell *i* is characterized by the tuning curve 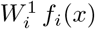 - the weights thus acting as gain modulations. For the analysis whose results are shown in Fig. 3 panels (C) and (D), we compute the Fisher information *F*_*code*_ making use of these effective tuning curves.

#### Optimal performances

To obtain the optimal performances the network model can achieve, we first consider a proxy network obtained by replacing the decision attractor network by a decision function *p*. This function results from a fit of the relation between the input received by the attractor network and the probability of choosing the most probable category.

This analog system is defined by the probability for the attractor network to respond correctly when a specific current is sent. We numerically obtain this probability by averaging the behavioral results over 1000 trials for a few different currents. Next we fit the resulting non-parametric function by analytic function. Given the observed shape of the empirical function, we fit with a sigmoid curve. This gives a decision function *p* that directly describes the behavior of the attractor network when it receives a specific current.

Given this function *p*, we then define the cost function as an average over stimulus space:

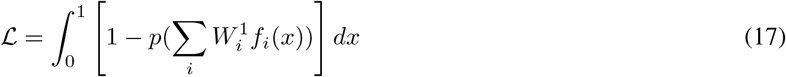

with *f*_*i*_(*x*) the tuning curve of neuron *i, x* being the stimulus input. We find the set of weights minimizing this cost function by solving a non-linear convex optimization problem, with the same set of constraints as in the neural circuit model: symmetry 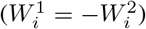, fixed norm 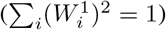, and 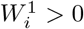.

In Fig. 5, Main Text, we compare the performances of the the network with this set of weights to the one obtained making use of the confidence-controlled Hebbian rule. Note that, since the optimal performances are here defined as an average over the ensemble of stimuli, a given learning algorithm may give better results on parts of the stimulus space. This is indeed the case for the CCHL, as can be seen in Fig. 5.

#### Confidence-controlled Hebbian learning as a stochastic gradient ascent algorithm

In this section, we exhibit the link between the confidence-modulated learning algorithm that we propose in this paper, and a gradient ascent algorithm derived within a machine learning approach. We consider the network obtained by replacing the attractor decision network by a softmax decision function. For this simpler network, we perform analytical computations in order to find the optimal update term to obtain maximal performances.

The network softmax output, giving the probability *P* (1|**r**) that the output neuron chooses category 1, is defined by:

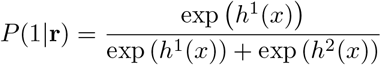

where, for *j* = 1, 2,

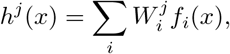

and *f*_*i*_(*x*) is the mean of the neural activity *r*_*i*_ of coding cell *i* when the input stimulus is *x*,

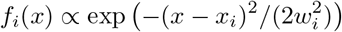

where *x*_*i*_ and *w*_*i*_ are the tuning curve parameters of cell *i*.

For simplicity, we impose for every 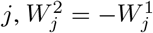, so that we also have *h*^2^(*x*) = −*h*^1^(*x*). Note that this is not essential, one can make the analysis without imposing the symmetry, with 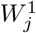 and 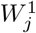. With the symmetry, the computation is easier to follow.

The goal of the network is here to maximize the reward averaged over all trials - *not* to estimate the correct probability *P* (1|*h*^1^(*x*)). We thus consider the following cost function (rather the reward function, which has to be maximized):

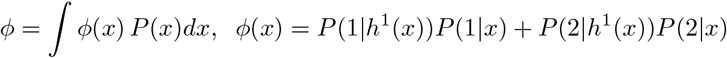

Up to a constant, we can rewrite

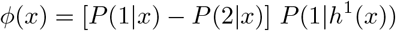

This cost is obviously maximum for *P* (1|*h*^1^(*x*)) = 1 if *P* (1|*x*) *> P* (2|*x*) and *P* (1|*h*^1^(*x*)) = 0 otherwise. Taking the derivative with respect to a weight 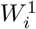, we get, for a given stimulus *x*, the updating rule

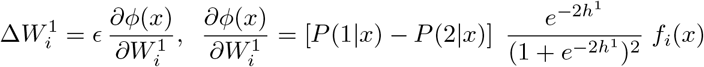

which we can rewrite as

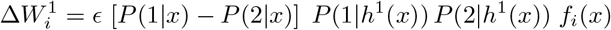

One recognize a Hebbian type updating rule (with the product of the input rate, *f*_*i*_, by the desired output - a graded form, *P* (1|*x*) −*P* (2|*x*)), with a modulation by the term *P* (1|*h*^1^(*x*)) *P* (2|*h*^1^(*x*)) which is significantly non zero only near the (current estimate of) the class boundary. This is thus quite similar to the Confidence controlled Hebbian rule.

In order to mimic the weight normalization in the biophysical model, we add a weight normalization constraint as follows:

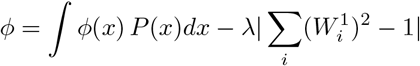

The gradient ascent updating rule is now:

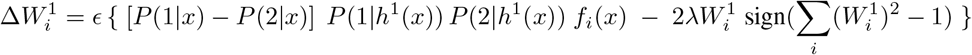

Now consider the case of ‘easy’ stimuli, that is such that the stimulus ambiguity is low - far from the category boundary. There, *P* (1|*h*^1^(*x*))*P* (2 |*h*^1^(*x*)) ≪ 1. This implies that for non ambiguous stimuli, the term coming from *ϕ*(*x*) is small compared to the one coming from the constraint. Hence the gradient is negative, making the weights to decrease. For neurons with preferred stimulus close to the boundary, *f*_*i*_(*x*) is equal to zero and the weights will be modified according to the normalization constraint alone. Hence, all the synaptic weights are decreasing. However, the decrease for the neurons with preferred stimulus close to the boundary is smaller than what it would have been in the absence of the term *P* (1|*h*^1^(*x*)) *P* (2|*h*^1^(*x*)).

If on the contrary the stimulus *x* is near the categories boundary, the term *P* (1|*h*^1^(*x*))*P* (2|*h*^1^(*x*))1 is close to 1*/*4. This implies that in that case the weights modification will always follow the sign of [*P* (1|*x*) −*P* (2|*x*)] for neurons close to the boundary. For tuning curves far from the boundary, *f*_*i*_(*x*) is close to 0 and the corresponding weights will decrease according to the normalization constraint.

The term *P* (1|*h*^1^(*x*))*P* (2|*h*^1^(*x*)) thus acts as a threshold on the weight modification. The modification coming from the cost *φ*(*x*) becomes negligible at some distance from the class boundary. This is quite similar to what does the Confidence controlled Hebbian rule. In our model with an attractor decision network, this effect is obtained making use of local variables provided by the network decision dynamics.

In the case of the standard reward-modulated Hebbian learning rule, the weight modification is, in average, of the form [*P* (1|*x*) −*P* (2|*x*)] *f*_*i*_(*x*) with an additional weights normalization term. Thus the term controlling the strength of the modification according to the distance to the boundary is not present.

## Data and Software availability

Software was written in the Julia (https://julialang.org/) programming language. Implementations of algorithms used to compute quantities presented in this study will be available at https://github.com/berlemontkevin/confidence_hebbian_learning.

## Supplementary Information

**Figure 6:**
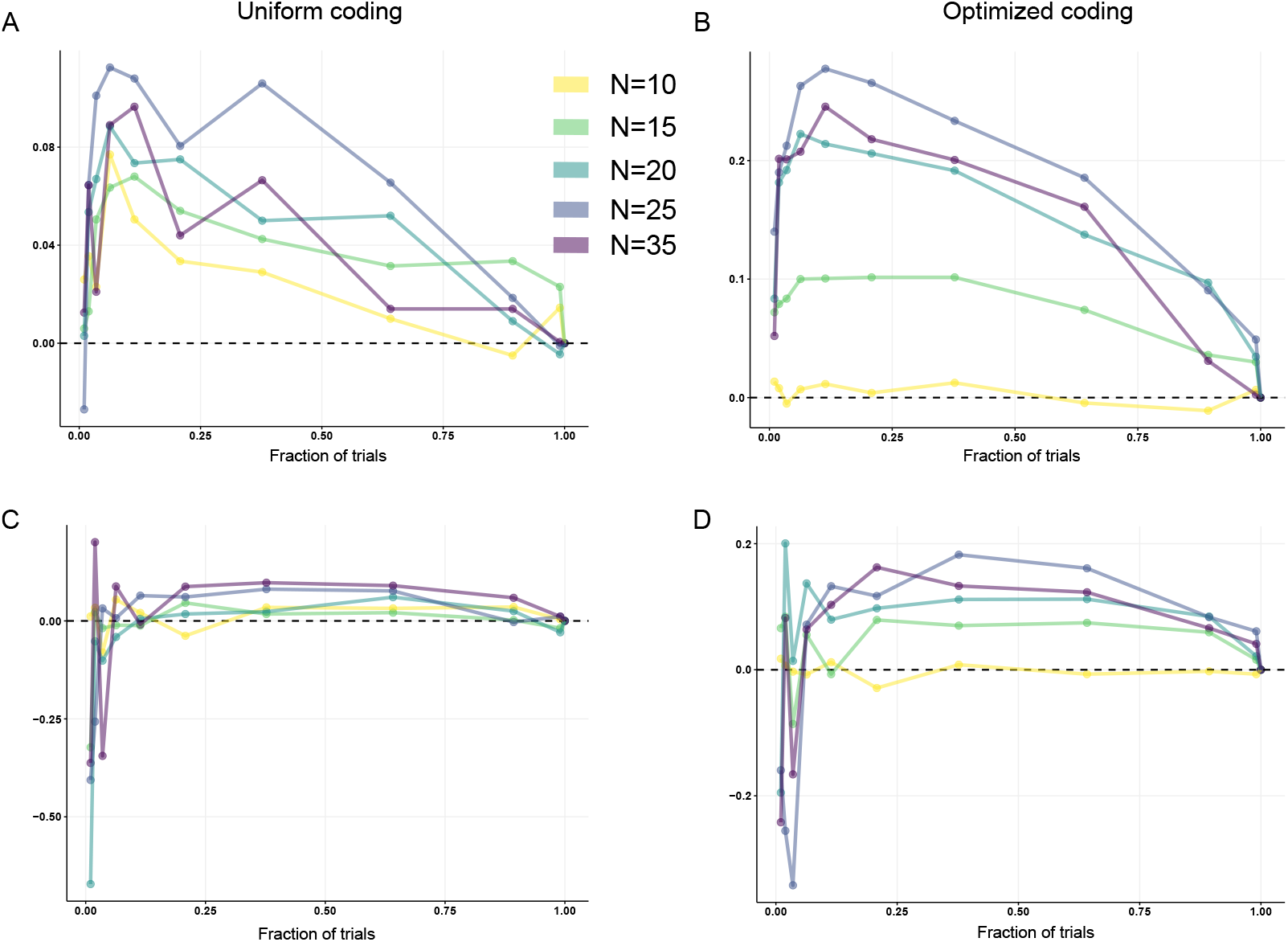
Influence of confidence on the learning process. (A): Effect of confidence modulation on the performances for a network with an optimized coding layer after 10000 trials and α = 0.25. The y-axis represents the difference between the performances, at an ambiguity x = 0.45, for a learning with modulation by confidence and for a learning without. The x-axis represents the percentage of trials where Δr was lower than the threshold z. (B): Same as (A) but for a neural circuit model with an uniformly distributed coding neurons. (C): Effect of confidence modulation on the performances for a network with an optimized coding layer after 1000 trials and α = 0.25. The y-axis represents the difference between the performances, at an ambiguity x = 0.45, for a learning with modulation by confidence and for a learning without. The x-axis represents the percentage of trials where Δr was lower than the threshold z. (D): Same as (C) but for a neural circuit model with an uniformly distributed coding neurons.

**Figure 7:**
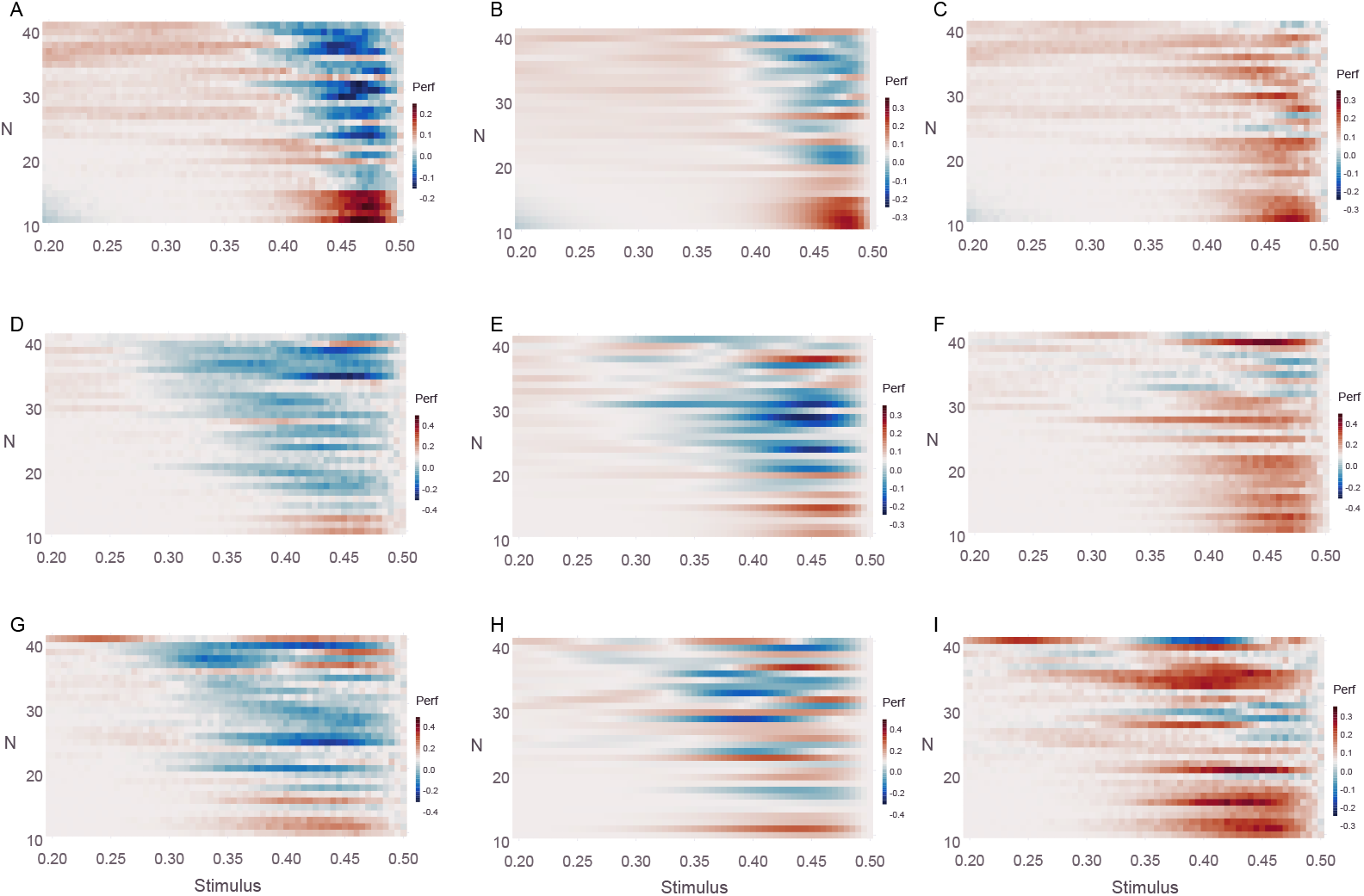
Late stage of learning. Difference of performances after learning (10000 trials) between networks with an uniform (or optimized) coding layer. Red (resp. Blue) corresponds to a positive (resp. negative) difference meaning that the network with the optimized coding layer has better (resp. worse) accuracy that the one with uniform layer. The x-axis corresponds to a variation of stimulus x, and the y-axis to the number of neurons N. (A), (D) and (G) correspond to the pure Hebbian learning rule described in the main text. The width of the categories is 0.2 for (A), 0.3 for (D) and 0.4 for (G). (B), (E) and (H) correspond to the reward modulated learning rule. The width of the categories is 0.2 for (B), 0.3 for (E) and 0.4 for (H). (C), (F) and (I) correspond to a learning with reward modulation by the confidence, with a threshold z = 15 Hz. The width of the categories is 0.2 for (C), 0.3 for (F) and 0.4 for (I).

**Figure 8:**
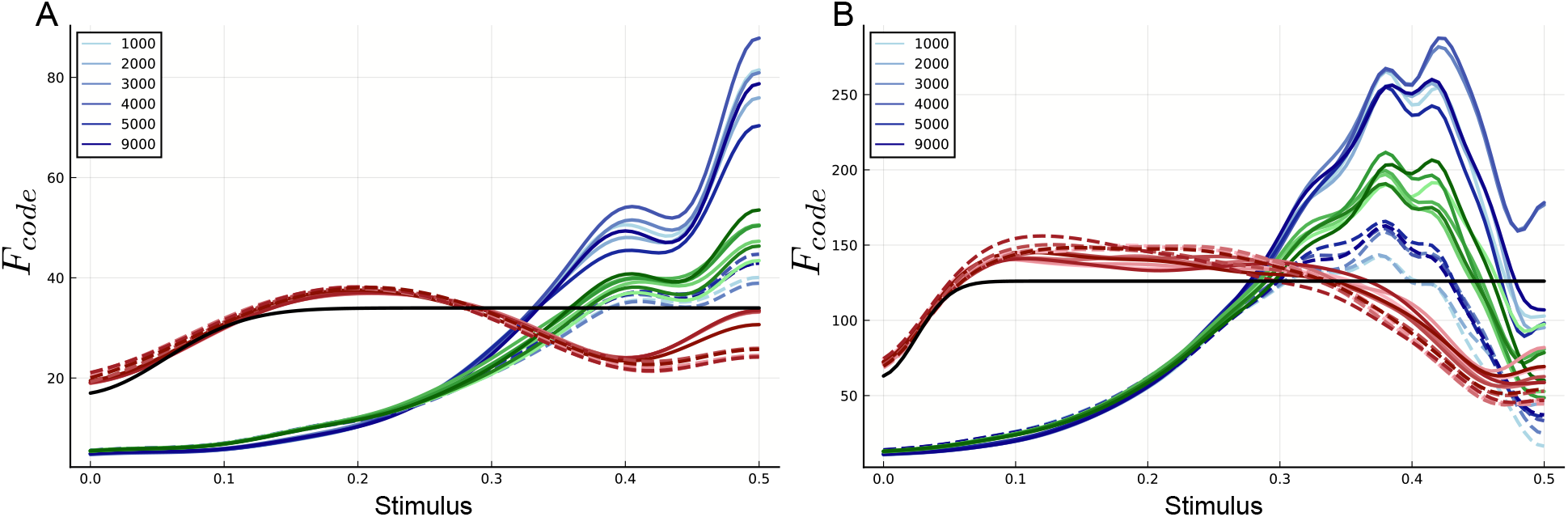
Fisher information of the coding layer taking into account the gain modulation due to the synaptic weights at late-stage learning, for N = 15, panel (A), and N = 35, panel (B). The gradient of color represents different snapshots of the network during learning (after an increasing number of trials, from 1000 to 9000). The network with an uniform coding layer is represented in red and the one with an optimized coding layer in blue. The dashed lines stands for the learning algorithm without confidence control, and the plain lines with the confidence control. The green color represents the optimized network with a reward-modulated learning rule. The width of the Gaussian categories is α = 0.25, the threshold for confidence controls is z = 15 Hz.

**Figure 9:**
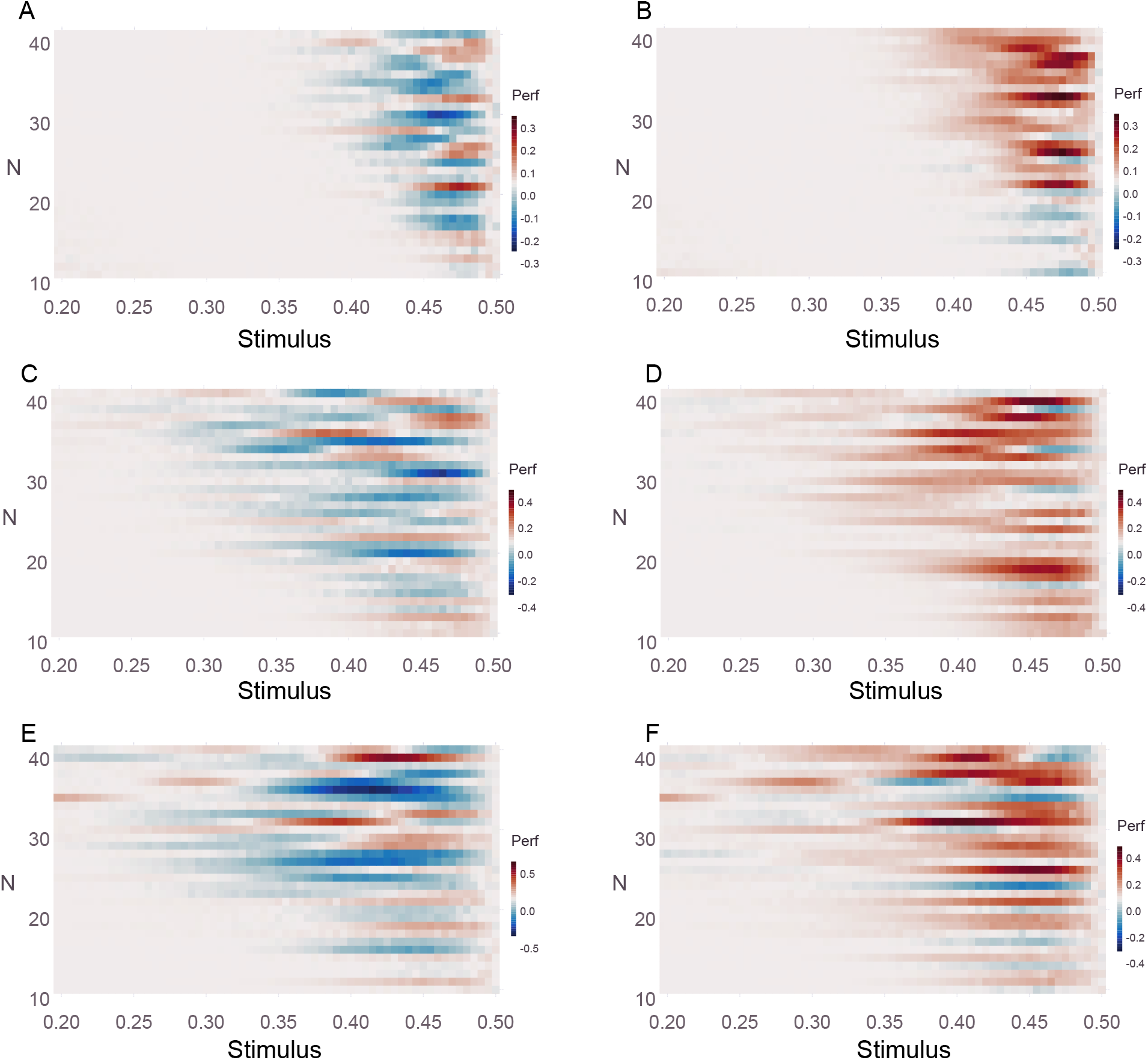
(A), (C), (E): Difference between reward-modulated Hebbian learning and pure Hebbian learning at α = 0.2 (A), 0.3 (C) and 0.4 (E) and 10000 learning trials, for the task-optimized network. Red (resp. Blue) corresponds to a positive (resp. negative) difference meaning that the performances for the neural network with pure Hebbian learning are higher (resp. less) than the ones with the standard reward-modulated. (B), (D), (F): Difference between reward-modulated Hebbian learning and confidence-controlled Hebbian learning at α = 0.2 (B), 0.3 (D) and 0.4 (F) and 10000 learning trials, for the task-optimized network. Red (resp. Blue) corresponds to a positive (resp. negative) difference meaning that the performances for the neural network with confidence-controlled Hebbian learning are higher (resp. less) than the ones with the standard reward-modulated.

**Figure 10:**
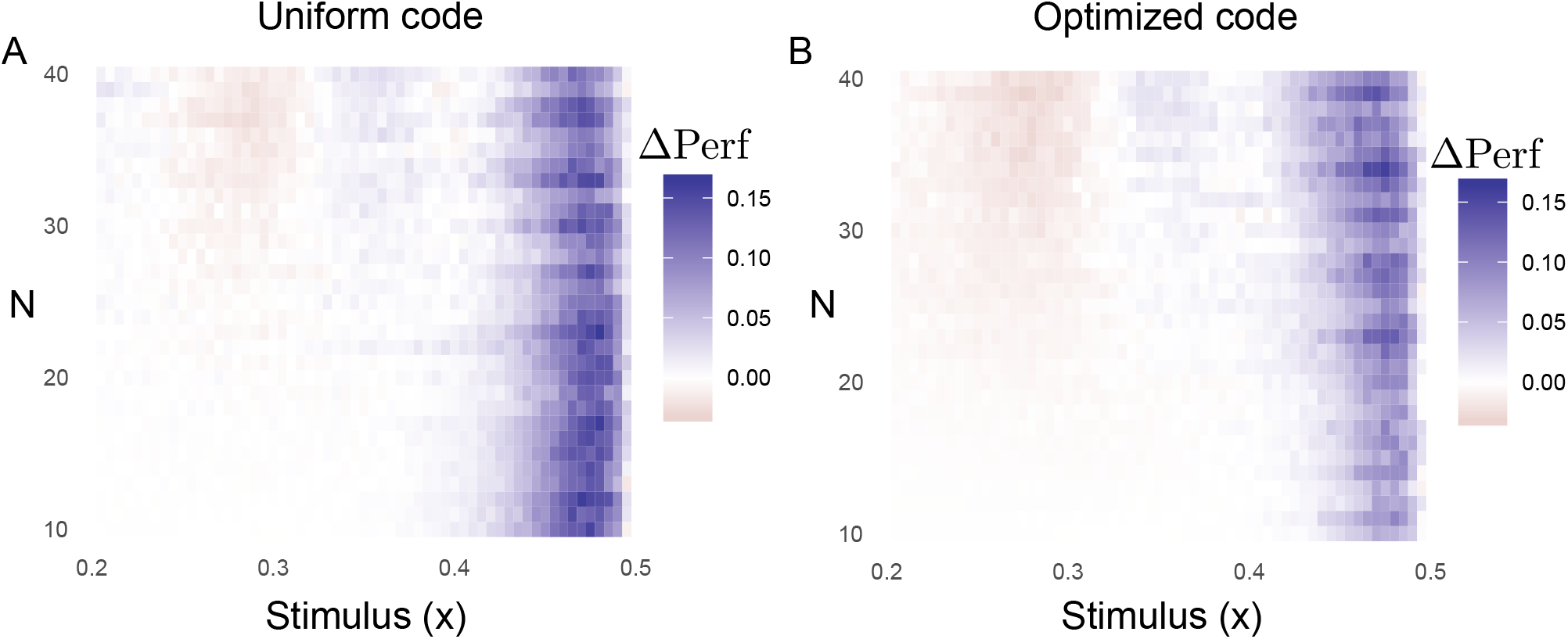
Difference between confidence-controlled Hebbian learning and optimal performances. (A) and (B) results for 10000 learning trials. Blue (resp. Red) corresponds to a positive (resp. negative) difference, meaning that the performances for the neural network with confidence-controlled learning are less (resp. higher) than the optimum performances for this neural architecture. (A) corresponds to the case where the coding layer is uniform, and (B) when it is optimized. All panels: categories of width α = 0.3.

## References

L. Abbott and F. S. Chance. Drivers and modulators from push-pull and balanced synaptic input. Progress in brain research, 149:147–155, 2005.

A. Alemi, C. Baldassi, N. Brunel, and R. Zecchina. A three-threshold learning rule approaches the maximal capacity of recurrent neural networks. PLoS computational biology, 11(8), 2015.

J. Y. Angela and P. Dayan. Uncertainty, neuromodulation, and attention. Neuron, 46(4):681–692, 2005.

F. G. Ashby and W. T. Maddox. Human category learning. Annu. Rev. Psychol., 56:149–178, 2005.

J. M. Beck, W. J. Ma, R. Kiani, T. Hanks, A. K. Churchland, J. Roitman, M. N. Shadlen, P. E. Latham, and A. Pouget. Probabilistic population codes for bayesian decision making. Neuron, 60(6):1142–1152, 2008.

T. E. Behrens, M. W. Woolrich, M.E. Walton, and M.F. Rushworth. Learning the value of information in an uncertain world. Nature neuroscience, 10(9):1214–1221, 2007.

K. Berlemont and J.-P. Nadal. Perceptual decision-making: Biases in post-error reaction times explained by attractor network dynamics. Journal of Neuroscience, 39(5):833–853, 2019.

K. Berlemont, J.-R. Martin, J. Sackur, and J.-P. Nadal. Does nonlinear neural network dynamics explain human confidence in a sequence of perceptual decisions? BioRxiv, page 648022, 2019.

K. Berlemont, J.-R. Martin, J. Sackur, and J.-P. Nadal. Nonlinear neural network dynamics accounts for human confidence in a sequence of perceptual decisions. Scientific reports, 10(1):1–16, 2020.

R. Bogacz, E. Brown, J. Moehlis, P. Holmes, and J. D. Cohen. The physics of optimal decision making: a formal analysis of models of performance in two-alternative forced-choice tasks. Psychological review, 113(4):700, 2006.

L. Bonnasse-Gahot and J.-P. Nadal. Neural coding of categories: information efficiency and optimal population codes. Journal of computational neuroscience, 25(1):169–187, 2008.

L. Bonnasse-Gahot and J.-P. Nadal. Perception of categories: from coding efficiency to reaction times. Brain Research, 1434:47–61, 2012.

J. A. Cromer, J. E. Roy, and E. K. Miller. Representation of multiple, independent categories in the primate prefrontal cortex. Neuron, 66(5):796–807, 2010.

J. Drugowitsch, A. G. Mendonça, Z. F. Mainen, and A. Pouget. Learning optimal decisions with confidence. Proceedings of the National Academy of Sciences, 116(49):24872–24880, 2019.

G. Dutilh, J. Vandekerckhove, B. U. Forstmann, E. Keuleers, M. Brysbaert, and E.-J. Wagenmakers. Testing theories of post-error slowing. Attention, Perception, & Psychophysics, 74(2):454–465, 2012.

T. A. Engel, W. Chaisangmongkon, D. J. Freedman, and X.-J. Wang. Choice-correlated activity fluctuations underlie learning of neuronal category representation. Nature communications, 6(1):1–12, 2015.

A. Fanini and J. A. Assad. Direction selectivity of neurons in the macaque lateral intraparietal area. Journal of neuro-physiology, 101(1):289–305, 2009.

M. Faraji, K. Preuschoff, and W. Gerstner. Balancing new against old information: the role of puzzlement surprise in learning. Neural computation, 30(1):34–83, 2018.

J. K. Fitzgerald, D. J. Freedman, and J. A. Assad. Generalized associative representations in parietal cortex. Nature neuroscience, 14(8):1075, 2011.

D. J. Freedman and J. A. Assad. Experience-dependent representation of visual categories in parietal cortex. Nature, 443 (7107):85–88, 2006.

N. Frémaux and W. Gerstner. Neuromodulated spike-timing-dependent plasticity, and theory of three-factor learning rules. Frontiers in neural circuits, 9:85, 2016.

J. B. Fritz, S. V. David, S. Radtke-Schuller, P. Yin, and S. A. Shamma. Adaptive, behaviorally gated, persistent encoding of task-relevant auditory information in ferret frontal cortex. Nature neuroscience, 13(8):1011, 2010.

D. Ganguli and E. P. Simoncelli. Efficient sensory encoding and bayesian inference with heterogeneous neural populations. Neural computation, 26(10):2103–2134, 2014.

W. Gerstner, M. Lehmann, V. Liakoni, D. Corneil, and J. Brea. Eligibility traces and plasticity on behavioral time scales: experimental support of neohebbian three-factor learning rules. Frontiers in neural circuits, 12:53, 2018.

G. M. Ghose, T. Yang, and J. H. Maunsell. Physiological correlates of perceptual learning in monkey v1 and v2. Journal of neurophysiology, 87(4):1867–1888, 2002.

J. I. Gold and M. N. Shadlen. The neural basis of decision making. Annual Review of Neuroscience, 30:535–574, 2007.

R. L. Goldstone. Influences of categorization on perceptual discrimination. Journal of Experimental Psychology: General, 123(2):178, 1994.

M. Guggenmos, G. Wilbertz, M. N. Hebart, and P. Sterzer. Mesolimbic confidence signals guide perceptual learning in the absence of external feedback. Elife, 5:e13388, 2016.

I. Guyon, N. Matic, V. Vapnik, et al. Discovering informative patterns and data cleaning., 1996.

S. Harnad. Categorical perception. 2003.

D. O. Hebb. The organization of behavior: a neuropsychological theory. J. Wiley; Chapman & Hall, 1949.

R. Herbrich, N. D. Lawrence, and M. Seeger. Fast sparse gaussian process methods: The informative vector machine. In Advances in neural information processing systems, pages 625–632, 2003.

S. A. Hillyard, K. C. Squires, J. W. Bauer, and P. H. Lindsay. Evoked potential correlates of auditory signal detection. Science, 172(3990):1357–1360, 1971.

J. Jaramillo, J. F. Mejias, and X.-J. Wang. Engagement of pulvino-cortical feedforward and feedback pathways in cognitive computations. Neuron, 101(2):321–336, 2019.

M. Jepma, P. R. Murphy, M. R. Nassar, M. Rangel-Gomez, M. Meeter, and S. Nieuwenhuis. Catecholaminergic regulation of learning rate in a dynamic environment. PLoS computational biology, 12(10), 2016.

R. Kiani and M. N. Shadlen. Representation of confidence associated with a decision by neurons in the parietal cortex. science, 324(5928):759–764, 2009.

H. Köver, K. Gill, Y.-T. L. Tseng, and S. Bao. Perceptual and neuronal boundary learned from higher-order stimulus probabilities. Journal of Neuroscience, 33(8):3699–3705, 2013.

W. Krauth and M. Mézard. Learning algorithms with optimal stability in neural networks. Journal of Physics A: Mathematical and General, 20(11):L745, 1987.

A. Lak, M. Okun, M. M. Moss, H. Gurnani, K. Farrell, M. J. Wells, C. B. Reddy, A. Kepecs, K. D. Harris, and M. Carandini. Dopaminergic and prefrontal basis of learning from sensory confidence and reward value. Neuron, 105(4): 700–711, 2020.

N. D. Lawrence and J. C. Platt. Learning to learn with the informative vector machine. In Proceedings of the twenty-first international conference on Machine learning, page 65, 2004.

R. Legenstein, D. Pecevski, and W. Maass. A learning theory for reward-modulated spike-timing-dependent plasticity with application to biofeedback. PLoS computational biology, 4(10), 2008.

R. Legenstein, S. M. Chase, A. B. Schwartz, and W. Maass. A reward-modulated hebbian learning rule can explain experimentally observed network reorganization in a brain control task. Journal of Neuroscience, 30(25):8400–8410, 2010.

Y. Loewenstein and H. S. Seung. Operant matching is a generic outcome of synaptic plasticity based on the covariance between reward and neural activity. Proceedings of the National Academy of Sciences, 103(41):15224–15229, 2006.

W. J. Ma, J. M. Beck, P. E. Latham, and A. Pouget. Bayesian inference with probabilistic population codes. Nature neuroscience, 9(11):1432–1438, 2006.

F. Meyniel. Brain dynamics for confidence-weighted learning. bioRxiv, page 769315, 2019.

F. Meyniel and S. Dehaene. Brain networks for confidence weighting and hierarchical inference during probabilistic learning. Proceedings of the National Academy of Sciences, 114(19):E3859–E3868, 2017.

K. D. Miller and D. J. MacKay. The role of constraints in hebbian learning. Neural computation, 6(1):100–126, 1994.

B. Min, D. P. Bliss, A. Sarma, D. J. Freedman, and X.-J. Wang. A neural circuit mechanism of categorical perception: top-down signaling in the primate cortex. bioRxiv, 2020.

M. R. Nassar, R. C. Wilson, B. Heasly, and J. I. Gold. An approximately bayesian delta-rule model explains the dynamics of belief updating in a changing environment. Journal of Neuroscience, 30(37):12366–12378, 2010.

M. R. Nassar, R. Bruckner, and M. J. Frank. Statistical context dictates the relationship between feedback-related eeg signals and learning. eLife, 8, 2019.

T. Ott, P. Masset, and A. Kepecs. The neurobiology of confidence: From beliefs to neurons. Cold Spring Harbor Symposia on Quantitative Biology, LXXXIII:9–16, 2019.

C. Ranganath and G. Rainer. Neural mechanisms for detecting and remembering novel events. Nature Reviews Neuroscience, 4(3):193–202, 2003.

R. Ratcliff. A theory of memory retrieval. Psychological review, 85(2):59, 1978.

W. Schultz. Predictive reward signal of dopamine neurons. Journal of neurophysiology, 80(1):1–27, 1998.

W. Schultz, P. Dayan, and P. R. Montague. A neural substrate of prediction and reward. Science, 275(5306):1593–1599, 1997.

N. Sigala and N. K. Logothetis. Visual categorization shapes feature selectivity in the primate temporal cortex. Nature, 415(6869):318–320, 2002.

C. Summerfield and F. P. De Lange. Expectation in perceptual decision making: neural and computational mechanisms. Nature Reviews Neuroscience, 15(11):745–756, 2014.

R. S. Sutton and A. G. Barto. Reinforcement learning: An introduction. MIT press, 2018.

C. I. Tajima, S. Tajima, K. Koida, H. Komatsu, K. Aihara, and H. Suzuki. Population code dynamics in categorical perception. Scientific reports, 6:22536, 2016.

S. Tajima, K. Koida, C. I. Tajima, H. Suzuki, K. Aihara, and H. Komatsu. Task-dependent recurrent dynamics in visual cortex. eLife, 6:e26868, 2017.

D. J. Tolhurst, J. A. Movshon, and A. F. Dean. The statistical reliability of signals in single neurons in cat and monkey visual cortex. Vision research, 23(8):775–785, 1983.

R. Varrier, H. Stuke, M. Guggenmos, and P. Sterzer. Sustained effects of corrupted feedback on perceptual inference. Scientific reports, 9(1):1–12, 2019.

X.-J. Wang. Probabilistic decision making by slow reverberation in cortical circuits. Neuron, 36(5):955–968, 2002.

Z. Wei and X.-J. Wang. Confidence estimation as a stochastic process in a neurodynamical system of decision making. Journal of neurophysiology, 114(1):99–113, 2015.

J. C. Whittington and R. Bogacz. Theories of error back-propagation in the brain. Trends in cognitive sciences, 23(3): 235–250, 2019.

K.-F. Wong and X.-J. Wang. A recurrent network mechanism of time integration in perceptual decisions. Journal of Neuroscience, 26(4):1314–1328, 2006.

Y. Xin, L. Zhong, Y. Zhang, T. Zhou, J. Pan, and N.-l. Xu. Sensory-to-category transformation via dynamic reorganization of ensemble structures in mouse auditory cortex. Neuron, 103(5):909–921, 2019.

H. A. Xu, A. Modirshanechi, M. P. Lehmann, W. Gerstner, and M. H. Herzog. Novelty is not surprise: Exploration and learning in human sequential decision-making. bioRxiv, 2020.

T. Yang and J. H. Maunsell. The effect of perceptual learning on neuronal responses in monkey visual area v4. Journal of Neuroscience, 24(7):1617–1626, 2004.

